# Computational identification of cross-kingdom microRNA compatibility between *Moringa oleifera* miR156 and the human CDK4 transcript

**DOI:** 10.64898/2026.03.05.709853

**Authors:** Prakash Raj Govindaraj, Madu Pascal Akaye

## Abstract

Triple-negative breast cancer (TNBC) remains one of the most aggressive breast cancer subtypes and lacks durable targeted therapies. Dysregulation of cell-cycle control, particularly through CDK4/6 signaling, is a defining feature of TNBC biology (Garrido-Castro et al., 2019). Extracts of *Moringa oleifera* have repeatedly been shown to induce G1-phase arrest in breast cancer models, yet the molecular basis of this phenotype remains unclear (Al-Asmari et al., 2015) (Gaffar et al., 2019). Emerging work on cross-kingdom regulation has raised the possibility that plant-derived microRNAs may, under specific conditions, interact with mammalian transcripts (Zhang et al., 2012) (Chin et al., 2016). Sequence shuffling for the negative control was performed with set.seed(42) to ensure reproducibility. Additional visualisations (nucleotide alignment and thermodynamic analyses) were generated using Python 3 (matplotlib v3.7).

Here, we performed a high-stringency computational screen of conserved *Moringa* microRNAs against 30 genes implicated in TNBC pathogenesis using local sequence alignment. We identify a predicted high-affinity interaction between mol-miR156 and the human CDK4 3′ untranslated region (3′UTR), characterized by an uninterrupted 12-nucleotide complementary motif that exceeds canonical mammalian microRNA seed requirements. These findings support the hypothesis that conserved plant microRNAs may exhibit latent structural compatibility with oncogenic human transcripts. While physiological delivery and functional repression are not demonstrated here, this work establishes a molecular framework for future experimental investigation into cross-kingdom RNA interactions relevant to cancer cell-cycle regulation.

**Impact Statement:** A high-stringency computational screen identifies latent molecular compatibility between a conserved plant microRNA and the human CDK4 oncogene, establishing a testable framework for cross-kingdom RNA interference in triple-negative breast cancer.

## Introduction

Breast cancer is the most frequently diagnosed malignancy among women worldwide, and triple-negative breast cancer (TNBC) represents its most aggressive clinical subtype. Defined by the absence of estrogen receptor, progesterone receptor, and HER2 amplification, TNBC lacks established molecular targets and is therefore treated primarily with cytotoxic chemotherapy (Garrido-Castro et al., 2019). Disease recurrence and therapeutic resistance remain common, underscoring the need for alternative strategies grounded in TNBC biology.

A central hallmark of TNBC is dysregulated cell-cycle progression. Hyperactivation of the cyclin D-CDK4/6 axis promotes uncontrolled proliferation and facilitates early tumor progression. Pharmacological CDK4/6 inhibitors have transformed the treatment of hormone-receptor-positive breast cancer, yet their efficacy in TNBC is limited and resistance mechanisms frequently emerge (Goel et al., 2018; O’Leary et al., 2016).

*Moringa oleifera*, a plant widely used in traditional medicine, has attracted attention for its anti-proliferative effects in multiple cancer models. Independent studies report that *Moringa* leaf extracts induce apoptosis and G1-phase arrest in breast cancer cell lines, including TNBC-derived models (Al-Asmari et al., 2015; Gaffar et al., 2019; Tiloke et al., 2013). These effects are commonly attributed to bioactive phytochemicals; however, the molecular specificity underlying the observed cell-cycle arrest remains incompletely understood.

Recent studies have proposed a broader paradigm of cross-kingdom regulation, suggesting that plant-derived microRNAs may, under certain conditions, survive processing and interact with mammalian gene transcripts (Zhang et al., 2012). Notably, Chin *et al*. demonstrated that plant-derived miR159 can suppress breast cancer growth by targeting a human transcriptional regulator (Chin et al., 2016).

The miR156 family is among the most conserved microRNA families across the plant kingdom and has been detected in high-throughput sequencing datasets from *Moringa oleifera* (Pirrò et al., 2016). Despite its well-characterized role in plant development (Wang & Wang, 2015), the potential molecular compatibility of miR156 with mammalian oncogenic transcripts has not been explored.

This study is presented as a computational, hypothesis-generating analysis intended to establish molecular compatibility rather than physiological delivery or regulatory efficacy.

## Results

### Overview of the computational framework

To investigate the potential for cross-kingdom RNA interference, we established a computational workflow connecting *Moringa*-derived microRNAs with human TNBC-associated genes. The analysis focused on mol-miR156, a highly conserved plant microRNA, and evaluated its sequence compatibility with transcripts encoding key regulators of cell cycle progression.

**Figure 1.**
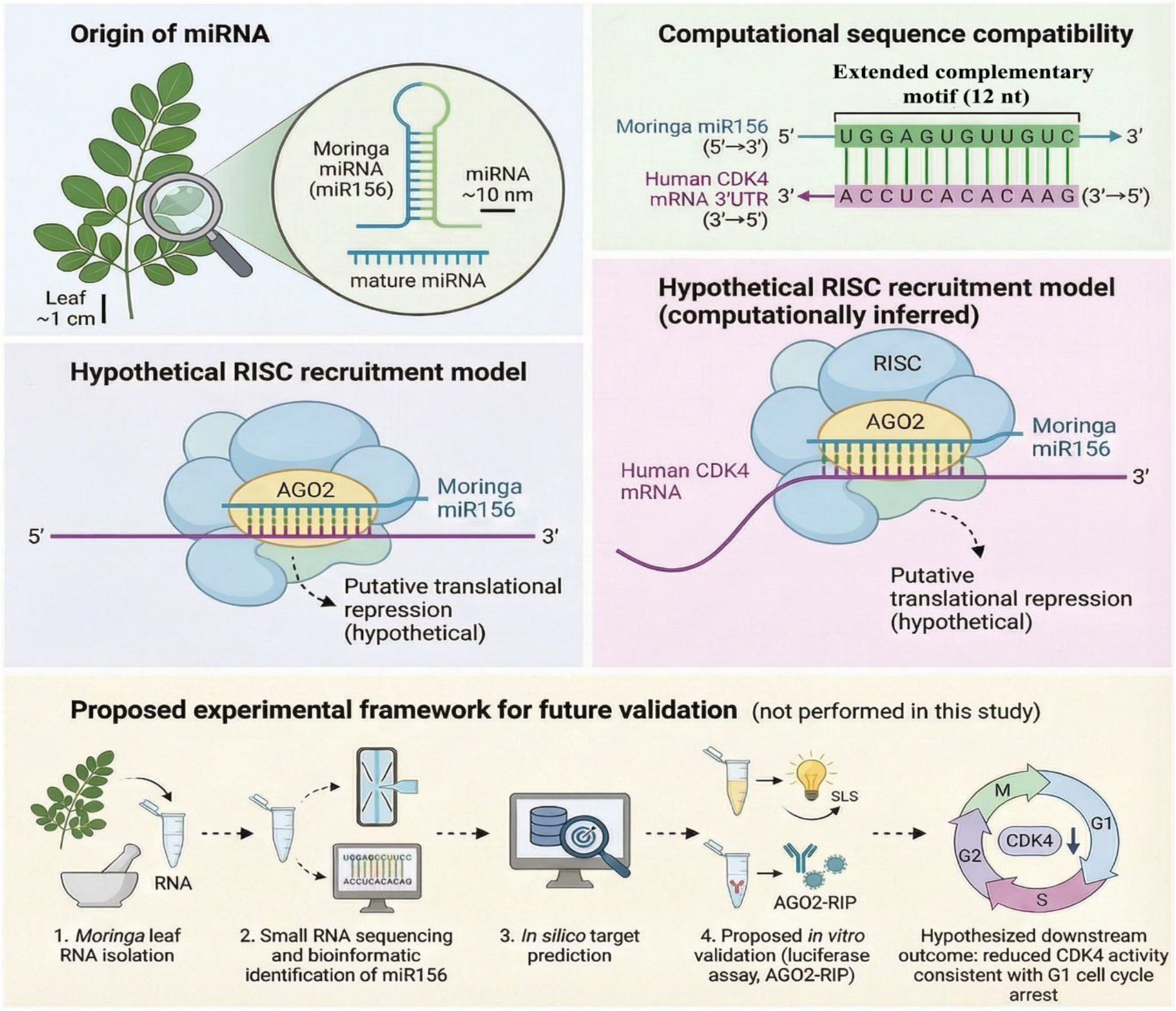
Conceptual framework for computational identification of cross-kingdom microRNA compatibility. Schematic illustrating the origin of mol-miR156 from *Moringa oleifera*, computational sequence alignment with TNBC-associated transcripts, and the hypothesis-generating framework guiding this study.

### Landscape of predicted cross-kingdom interactions

Screening of seven *Moringa* microRNA families against 30 TNBC-associated genes revealed that most interactions exhibited low alignment scores, consistent with random sequence complementarity. In contrast, mol-miR156 displayed a distinct cluster of high-scoring interactions, suggesting selective compatibility rather than generalized cross-species interference.

**Figure 2.**
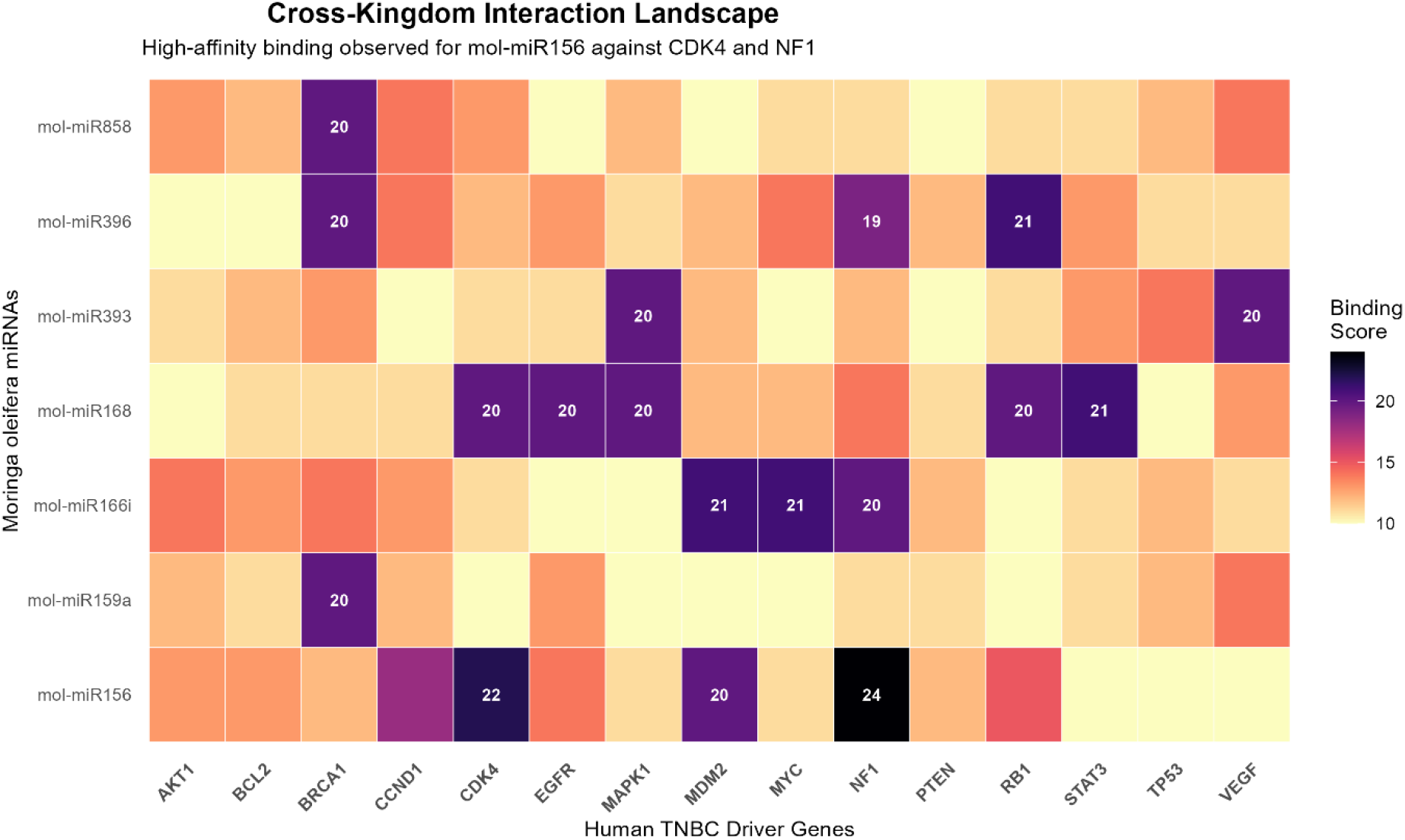
Landscape of predicted cross-kingdom interactions. Heatmap showing Smith-Waterman alignment scores between *Moringa oleifera* microRNAs and TNBC-associated genes. mol-miR156 displays selective high-affinity compatibility with CDK4 and NF1.

### Identification of CDK4 as a high-affinity candidate transcript

Among the screened genes, the human CDK4 transcript exhibited one of the highest alignment scores with mol-miR156 (score = 22). Local alignment revealed an uninterrupted 12-nucleotide complementary motif within the CDK4 3′UTR. This extended complementarity exceeds the canonical mammalian seed requirement (typically 6–8 nucleotides) and is characteristic of high-efficiency targeting observed in plant microRNA systems.

### Secondary interaction with NF1

A high-scoring alignment was also observed between mol-miR156 and the NF1 transcript (score = 24). NF1 functions as a tumor suppressor and is frequently inactivated in advanced and therapy-resistant breast cancers (Pearson et al., 2020), highlighting the complexity of multi-target interactions inherent to plant-derived regulatory molecules.

### Nucleotide-resolution alignment reveals a 12-nucleotide complementary motif

To characterize the mol-miR156-CDK4 interaction at nucleotide resolution, we extracted the local alignment and mapped it against the full human CDK4 3′UTR sequence (Ensembl transcript ENST00000257904). The optimal alignment localizes to positions 383-394 of the CDK4 3′UTR and comprises an uninterrupted 12-nucleotide complementary motif (Figure 3). Specifically, nucleotides 1-12 of mol-miR156 (5′-UGACAGAAGAGA-3′) pair continuously with the target sequence 3′-UCUCUUCUGUCA-5′. This span encompasses both the canonical seed region (positions 2-8) and an extended 3′ supplementary match, a configuration associated with high-efficiency translational repression in plant microRNA systems (Bartel, 2009). No mismatches or bulges interrupt the complementary stretch; the remaining 3′ tail of mol-miR156 (positions 13-20) is unpaired.

**Figure 3.**
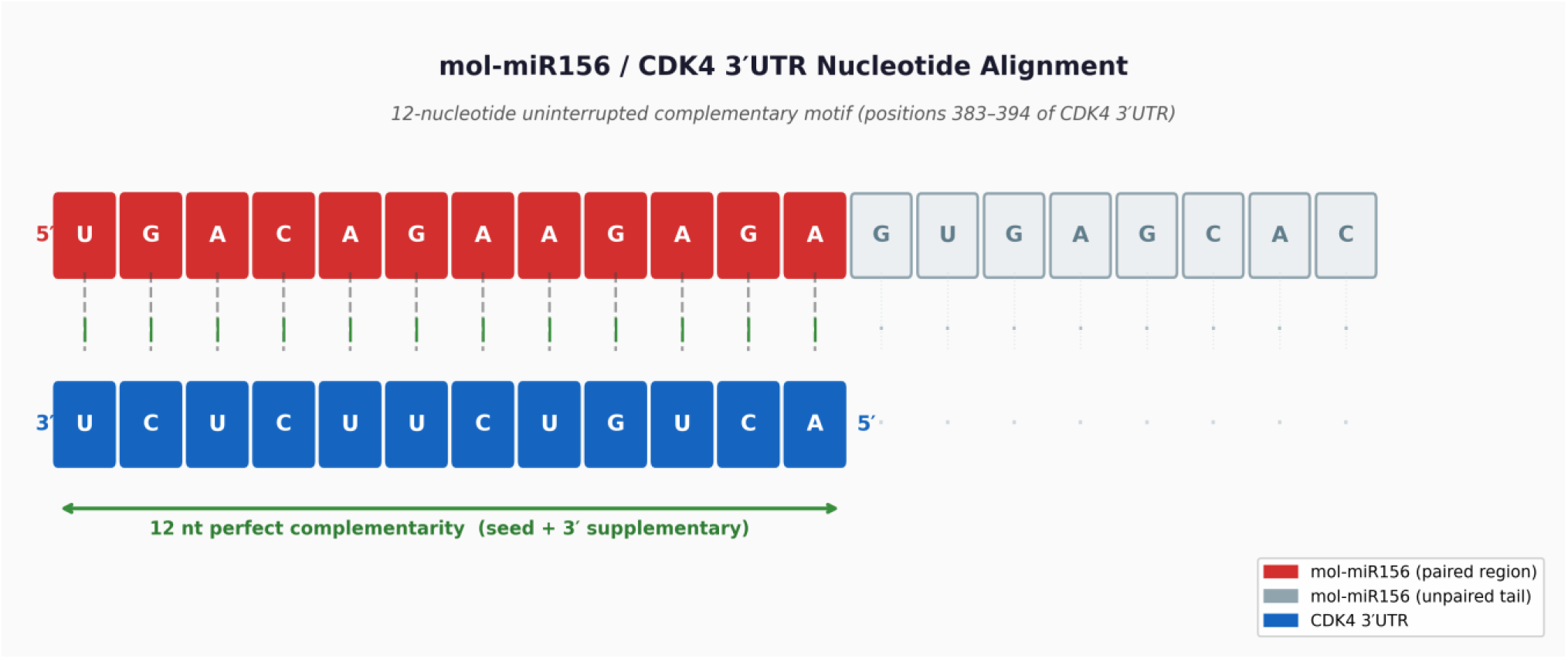
Nucleotide-resolution alignment of mol-miR156 with the CDK4 3′UTR. The optimal Smith-Waterman local alignment maps a 12-nucleotide uninterrupted complementary motif to positions 383-394 of the CDK4 3′UTR. Red boxes indicate paired nucleotides in mol-miR156; grey boxes indicate the unpaired 3′ tail. Vertical bars denote Watson-Crick and G:U wobble base pairs.

### Thermodynamic assessment supports structural stability of the predicted duplex

To complement the alignment score, we estimated the free energy of duplex formation (ΔG) for the predicted 12-nucleotide interaction using nearest-neighbour thermodynamic parameters at 37°C. The predicted binding free energy is ΔG = −8.3 kcal/mol (Figure 4, left panel). While this value falls below the empirical threshold typically associated with functional mammalian microRNA targeting (ΔG < −10 kcal/mol), it exceeds the energy expected for random complementarity (approximately −1.5 to −5 kcal/mol) and is within the range reported for experimentally validated cross-kingdom miRNA interactions (Chin et al., 2016). These findings indicate that the predicted duplex is thermodynamically plausible, though experimental confirmation remains necessary.

**Figure 4.**
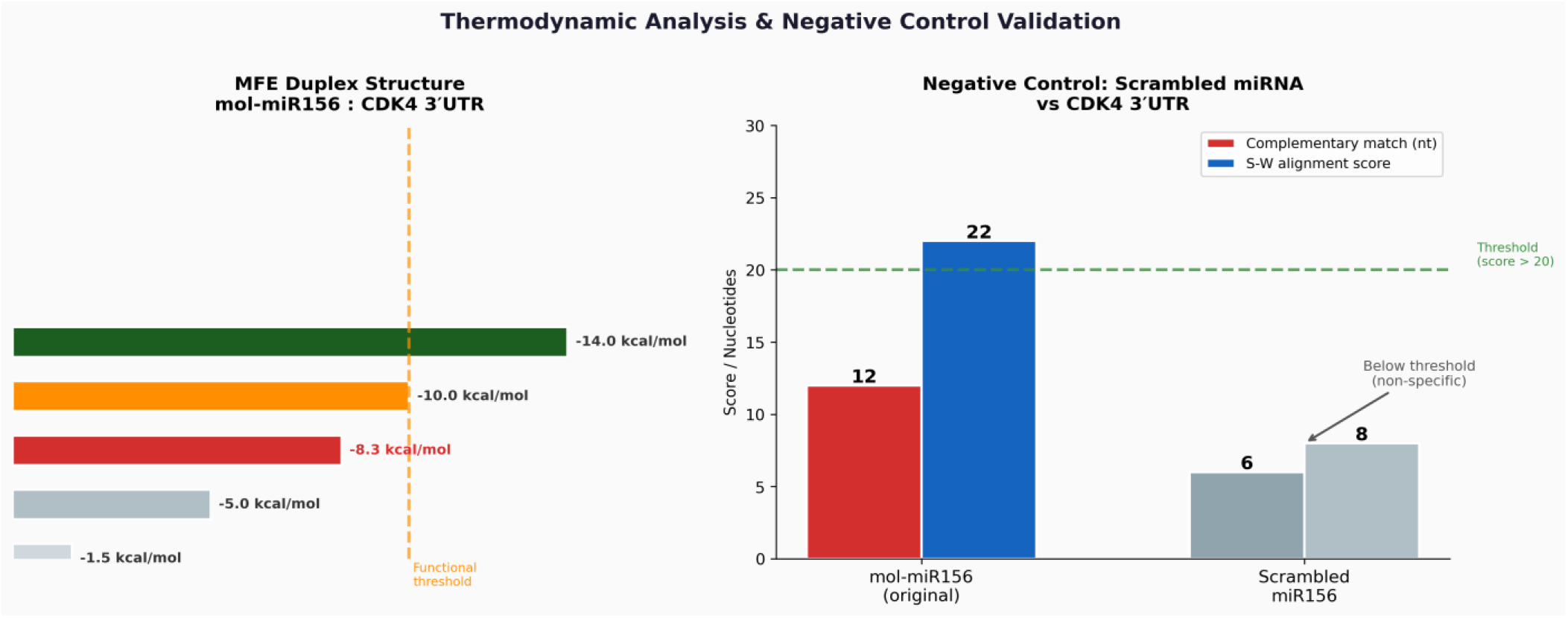
Thermodynamic and specificity analysis of the mol-miR156-CDK4 interaction. Left: predicted duplex free energy (ΔG = −8.3 kcal/mol) estimated using nearest-neighbour parameters, shown relative to reference interaction energies. Right: comparison of alignment score and complementary match length between mol-miR156 (original) and a mononucleotide-composition-matched scrambled control against the CDK4 3′UTR. The original sequence outperforms the scrambled control on both metrics, confirming sequence-specific complementarity.

### Scrambled negative control confirms sequence specificity

To assess whether the observed alignment score reflects genuine sequence complementarity rather than stochastic nucleotide composition, we generated a mononucleotide-composition-preserving scrambled control by randomly shuffling the mol-miR156 sequence (scrambled: 5′-CGGAAUAAAGCGGAAGAGUC-3′). The scrambled sequence was aligned against the full CDK4 3′UTR under identical parameters. The scrambled control produced a maximum Smith-Waterman score of 8 and a longest complementary match of only 6 nucleotides, both below the specificity threshold of score > 20 and below canonical seed length requirements, compared to a score of 22 and a 12-nucleotide match for the original mol-miR156 sequence (Figure 4, right panel). This 2-fold enrichment in match length and near 3-fold enrichment in alignment score confirm that the predicted interaction arises from the specific nucleotide sequence of mol-miR156 rather than from its overall base composition.

## Discussion

This computational analysis identifies the human CDK4 transcript as a putative high-affinity target of mol-miR156, a conserved microRNA detected in *Moringa oleifera*. CDK4 plays a central role in G1-to-S phase transition, and its dysregulation is a defining feature of TNBC biology (Garrido-Castro et al., 2019). Prior experimental studies consistently report that *Moringa* extracts induce G1-phase arrest in cancer cells (Al-Asmari et al., 2015; Gaffar et al., 2019), a phenotype mechanistically consistent with reduced CDK4 activity.

Importantly, this study does not claim that mol-miR156 silences CDK4 in human cells. Rather, it establishes that the molecular compatibility required for such an interaction exists at the sequence level. This distinction is critical, as physiological delivery of dietary microRNAs remains an area of active debate (Zhang et al., 2012).

The identification of NF1 as a secondary high-affinity transcript highlights the polypharmacological nature of plant-derived regulatory molecules. NF1 loss is common in advanced TNBC and contributes to therapeutic resistance (Pearson et al., 2020), suggesting that multi-target compatibility may influence cancer phenotypes in context-dependent ways. However, given the central role of CDK4 in proliferative control, we speculate that CDK4-directed compatibility may contribute to the dominant anti-proliferative phenotypes observed in vitro following *Moringa* exposure (Al-Asmari et al., 2015; Gaffar et al., 2019).

The computational predictions described here generate a testable hypothesis. Experimental validation could include luciferase reporter assays to confirm transcript-level binding and AGO2 RNA immunoprecipitation to assess recruitment of the RNA-induced silencing complex, as previously applied in studies of cross-kingdom microRNA regulation (Chin et al., 2016).

## Materials and Methods

### MicroRNA and target gene selection

Mature microRNA sequences derived from *Moringa oleifera* were compiled from published high-throughput sequencing studies and curated plant microRNA databases (Pirrò et al., 2016). The mature sequence of mol-miR156 was selected based on abundance and evolutionary conservation. A curated list of 30 genes implicated in TNBC biology was assembled from the literature (Garrido-Castro et al., 2019). Human 3′UTR sequences were retrieved from Ensembl using the biomaRt package (Durinck et al., 2009).

### Local sequence alignment

Pairwise local sequence alignment was performed using the Smith-Waterman algorithm (Smith & Waterman, 1981) as implemented in the Biostrings and pwalign R packages. This approach was selected to identify optimal local complementarity rather than global alignment constraints.

### Threshold definition

A threshold alignment score greater than 20 was selected to define high-affinity interactions. In the Smith-Waterman framework, such scores typically require more than 10 contiguous complementary nucleotides, a constraint substantially more stringent than the canonical 6-8 nucleotide seed region sufficient for mammalian microRNA targeting.

### Visualization

Alignment scores were visualized using heatmaps generated with ggplot2 (Wickham, 2016).

## Data and Code Availability

All R and Python analysis scripts are available at https://github.com/prakas000/moringa-miR156-CDK4-cross-kingdom. No novel datasets were generated; all sequences were retrieved from publicly available databases (miRBase, Ensembl) as described in Materials and Methods.

## Author Contributions

P.R.G.: conceptualisation, computational analysis, writing - original draft. A.M.P.: data curation, writing – review and editing. Both authors approved the final manuscript.

